# Toward Crosstalk-free All-optical Interrogation of Neural Circuits

**DOI:** 10.1101/2025.06.18.660335

**Authors:** Gewei Yan, Guangnan Tian, Yiming Fu, Yingzhu He, Zhentao She, Kenny K.Y. Chen, Julie L. Semmelhack, Jianan Y. Qu

## Abstract

All-optical interrogation, based on high-resolution two-photon stimulation and imaging, has emerged as a potentially transformative approach in neuroscience, allowing for the simultaneous precise manipulation and monitoring of neuronal activity across various model organisms. However, the unintended excitation of light-gated ion channels such as channelrhodopsin (ChR) during two-photon calcium imaging with genetically encoded calcium indicators (GECIs) introduces artifactual neuronal perturbation and contaminates neural activity measurements. In this study, we propose an active pixel power control (APPC) approach, which dynamically adjusts the imaging laser power at each scanning pixel, to address the challenge. We aim to achieve simultaneous two-photon optogenetic manipulation and calcium imaging with a single femtosecond laser, while minimizing the crosstalk between manipulation and imaging. To study this technology’s capabilities, we applied it to the larval zebrafish brain *in vivo.* Our results demonstrate that the APPC approach preserves GECI signal quality while suppressing optogenetic artifacts significantly. This enhances the accuracy of neural circuit dissection and advances the precision of all-optical interrogation, offering a robust framework for probing neural circuit dynamics and causality *in vivo* with high fidelity, potentially across various model organisms. Importantly, this technology can be seamlessly integrated with commonly used two-photon microscope systems in laboratories worldwide.

## INTRODUCTION

All-optical interrogation of neuronal circuits represents a paradigm-shifting advancement in neuroscience, successfully applied to various model organisms, including *Mus musculus* (mice)^1– 7^, *Danio rerio* (zebrafish)^8,9^, *Drosophila melanogaster* (fruit flies)^10^, and *Caenorhabditis elegans* (nematodes)^11,12^. This technique integrates genetically encoded calcium indicators (GECIs) with optogenetic actuators, such as channelrhodopsin (ChR), facilitating non-invasive, *in vivo* recording and manipulation of neuronal activity. Recent advancements in two-photon microscopy and holographic optogenetics enable neural ensemble recording and optogenetic control at single-cell resolution and sub-millisecond timescales^2,5,6,13,14^. Furthermore, closed-loop systems allow real-time extraction of neural activity to guide precise perturbations such as spike rates, timing, and synchrony^1,15,16^. These capabilities enable targeting of individual neurons based on functional signatures and advance mechanistic studies of circuit connectivity, plasticity, and disease-associated aberrant activity in conditions such as Alzheimer’s disease^17,18^ and Parkinson’s disease^19,20^. By elucidating neural circuit dynamics under targeted perturbation, this approach goes beyond merely correlating activity with behavior or cognitive states, establishing causal links; for example, it has been used to demonstrate how emotional states are encoded in neural attractor networks^2,21–25^. However, a major challenge emerges when a two-photon light source serves dual roles as both reader and stimulator in all-optical interrogation systems: crosstalk between these modules compromises experimental precision, and must be effectively minimized^3,6,14,26^.

Generally, there are three primary mechanisms underlying the crosstalk between calcium imaging and optogenetic stimulation^6,26,27^. First, functional imaging laser scanning may stimulate opsins to induce ectopic neural activity. Second, the stimulation laser can contaminate calcium signals, thereby introducing recording artifacts. Third, ChR expressed in densely packed neurons may lead to off-target activation during optogenetic stimulation. While existing strategies mitigate some issues, such as image-processing algorithms to correct calcium signal distortions^9,28,29^ and soma-localized opsin variants to restrict ChR expression to soma membranes^6,26^, the first issue, imaging-induced opsin activation, remains a general but unsolved problem. Though this first type of crosstalk could be avoided in single-photon systems due to GCaMP’s superior single-photon absorption compared to ChRs^30–32^, it is exacerbated in two-photon systems, where ChR2, the most commonly used light gated ion channel protein, exhibits 10-fold higher two-photon absorption cross-sections than GCaMP^33,34^, the most widely used GECI. This property enables ChR2 to be activated at <10% of the laser power required for GCaMP imaging^35^. Efforts to reduce crosstalk via spectral separation by using red-shifted GECIs (e.g., RCaMP^36^, jRGECO^33^) or ChRs (e.g., ReaChR^37^, C1V1^38^, Chrones^39^, Chrimson^39,^ and ChRmine^2^) face inherent challenges, as these reagents were developed using single-photon excitation spectra as benchmarks, while two-photon excitation operates via a different quantum mechanism compared to single-photon excitation. The pronounced difference in the two-photon excitation spectrum results from more complex transitions between energy states^40^. Although red-shifted ChRs minimize single-photon activation at 488 nm (e.g., ChrimsonR retains around 25% of maximal efficiency at this wavelength^39^), their two-photon excitation efficiency remains high (about 50% at 920 nm for ChrimsonR)^6^, overlapping with GCaMP’s optimal two-photon imaging wavelength range. While pairing red GECIs with blue ChRs at 1100 nm eliminates crosstalk^26,41^, current red indicators suffer from limited dynamic range and signal-to-noise ratios^33^. Moreover, dual femtosecond laser systems for spectral separation are cost-prohibitive and technically constrained, hindering widespread adoption in laboratories worldwide that are equipped with standard two-photon systems using a single excitation laser.

To address these limitations, we demonstrated a dynamic imaging power control strategy employing active and precise control of excitation power at each scanning pixel utilizing a fast acousto-optic modulator (AOM). We tuned down the imaging power at designated pixels to reduce crosstalk while maintaining the efficient extraction of calcium signals. Our method utilizes ChR expression maps generated by standard fluorescent labeling to create an “Active Pixel Power Control” (APPC) scheme that selectively attenuates laser intensity or even blocks the laser at ChR+ pixels during two-photon calcium imaging. Specifically, synchronized with resonant scanning, the AOM dynamically adjusts power in real time, minimizing unintended ChR activation while maintaining calcium signal fidelity through spatial summation of pixel fluorescence intensities. We validated this approach in the larval zebrafish, a vertebrate model ideal for whole-brain imaging and optogenetics. We used a single femtosecond laser to combine two-photon calcium imaging (via GCaMP) with holographic optogenetic stimulation via spatial light modulator (SLM), eliminating the need for costly and complicated dual-laser excitation systems. Calcium signals were deconvolved to estimate spike rates^1,37^, enabling crosstalk quantification through brain-wide activity patterns. The results demonstrated that the APPC technology achieves nearly crosstalk-free all-optical interrogation without relying on protein engineering or spectral separation, while concurrently enabling precise, high-speed *in vivo* calcium imaging. By controlling laser power at single-pixel resolution, this strategy preserved signal quality while offering a scalable and practical solution for studying neural circuits in behaving animals. This advance potentially opens new avenues for probing causality in brain-wide dynamics and accelerates therapeutic development for neurological disorders in any laboratory equipped with a standard two-photon microscope system.

## RESULTS

### All-optical interrogation system with precise pixel-specific excitation power control

To achieve active pixel power control (APPC), which provides dynamic, pixel-specific control of imaging laser power based on ChR expression patterns to minimize crosstalk, we developed a two-photon microscope system with a single excitation source and a custom beam control module (Figure 1a and Figure S1). The excitation source is a Ti:Sapphire femtosecond laser (Chameleon Ultra II, Coherent) that can be tuned to wavelengths optimal for imaging both GCaMP and the red fluorescent protein fused to the channelrhodopsin. A polarized beam splitter separates the laser beam into independent imaging and optogenetic stimulation paths. In the imaging path, an electrically tunable lens (ETL) with resonant scanners enables high-speed volumetric imaging, capturing a 10-plane brain volume (400 × 400 × 120 µm^3^) at 3 Hz. An AOM, synchronized with the resonant scanner, dynamically controls the excitation power, while a dispersion prism pair compensates for wavelength-dependent AOM-induced dispersion. The AOM exhibits a rise time (τ*_on_*) of 85 ns, compatible with a pixel dwell time of 180 ns, enabling single-pixel resolution modulation. Using the 0th-order beam (5-100% power range), we calibrated the relationship between AOM driver voltage and output power (Figure 1b). We then confirm the second-order relationship of excitation power and excitation intensity, which has a maximum excitation contrast ratio of ∼400:1(Figure 1c). After adjusting the time delay (see Methods and Figure S2), we applied APPC imaging to a uniform fluorescent plate, demonstrating robust continuous control of excitation power at designated pixels (Figure 1d). Next, we tested the APPC on larval zebrafish and verified its ability to efficiently turn off the two-photon excitation of specific target neurons (Figure 1e, f). When applied to live, large-volume brain imaging in larval zebrafish, the AOM turned off the scanning laser in the eye region during functional imaging sessions (Figure S3a), effectively mitigating laser-induced agitation (Figure S3b-d).

**Figure 1.**
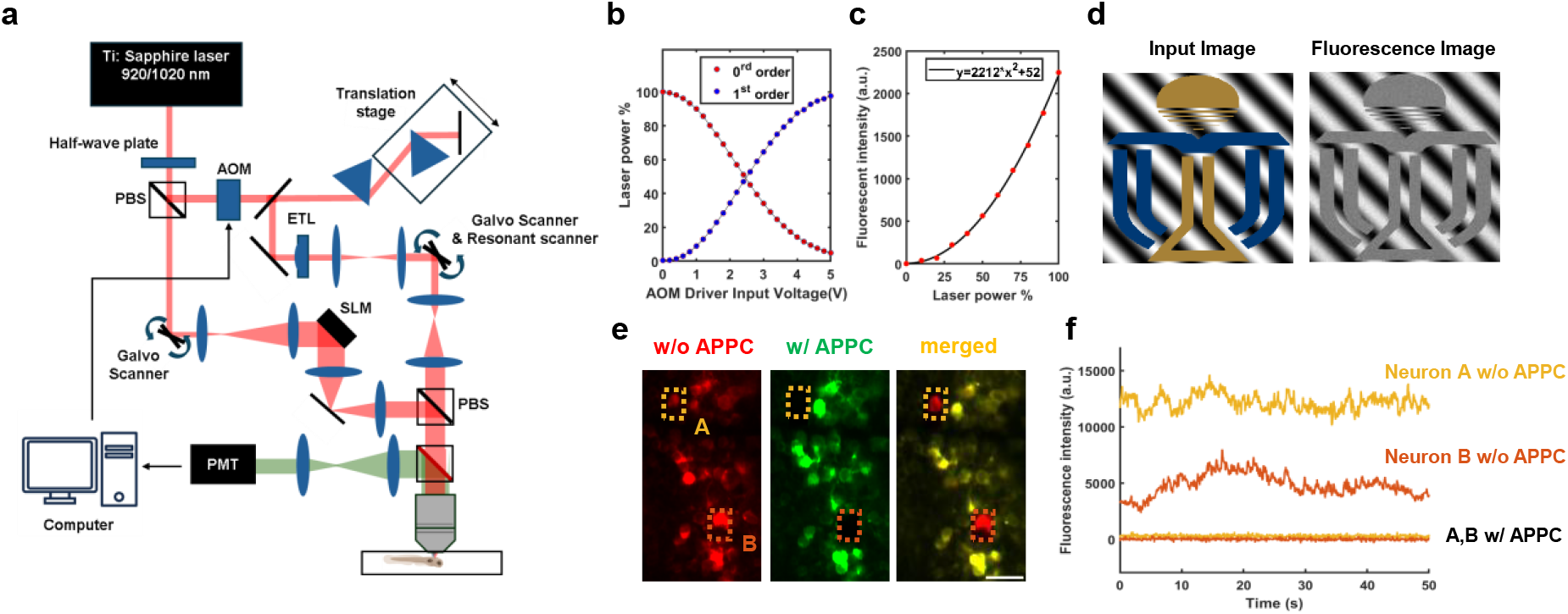
Two-photon imaging and stimulation microscope of active pixel power control (APPC). **a**, Simplified schematic representation of the two-photon optical interrogation system. (See Figure S1 for the complete version of the optical system.) **b**, AOM driver input voltage vs laser power. The 0^th^ order laser power can be continuously tuned between 5-100%, corresponding to a control voltage 5-0V. **c**, Fluorescent intensity vs laser power. The two-photon excited signal was measured by the fluorescence intensity of fluorescent beads, fitted by a second-order polynomial. **d**, APPC image of a fluorescent plate. Left: original input image. Right: two-photon fluorescence image of the fluorescent plate with the input image loaded on the AOM. **e**, Selective GCaMP6s stimulation of neurons via APPC in zebrafish larvae. Left: Calcium image (red) without APPC. Middle: Image (green) with APPC applied to block neuron A and B. Right: Merged image showing a comparison between conditions without and with APPC. **f**, Fluorescence trace of GCaMP for neurons A and B, with and without APPC.

In the optogenetics stimulation path, we programmed the SLM to generate scanning spot arrays targeting neuronal soma. Co-calibration of imaging and stimulation modules using fluorescent plates ensured spatial precision (see Methods).

### Crosstalk between imaging and stimulation for optogenetic targeted neurons

We first assessed the crosstalk of ChR2, the most widely used optogenetic tool across various animal models. Our strategy was to perform multi-plane volumetric imaging (3 Hz) under standard zebrafish imaging conditions^9,42^ while applying APPC to adjust laser intensity specifically in ChR2+ pretectal neurons. Next, we analyzed the calcium trace and evaluated pretectal neuron activity under different laser powers for the following APPC pattern updating (Figure 2a). In this study, we utilized a transgenic fish line: *Tg(KalTAu508; UAS-ChR2(H134R)-mCherry; UAS-GCaMP6)*^9,43,44^, in which the Gal4 variant protein KalTA4 drives expression of both ChR2-mCherry and GCaMP6s in specific neuron populations. Structural imaging confirmed colocalization of green (GCaMP6s) and red (mCherry-tagged ChR2) fluorescence in KalTA4u508+ pretectal neurons (Figure 2b). For calcium imaging, the laser power on pretectal neurons was set to 5 mW, 10 mW, and 15 mW, the commonly used imaging power for zebrafish^9^. We determined that 5 mW was the minimum laser power required to maintain an acceptable signal-to-noise ratio (SNR) for measuring calcium traces (ΔF/F) in individual ChR2+ neurons (Figure S4). Next, we performed three trials (100 s each) for every power condition, resulting in nine trials presented in a randomized sequence. Calcium traces (ΔF/F) from each neuron were normalized to their corresponding 15 mW trials (see Methods). Representative traces are shown in Figure 2c. Our results demonstrated that the normalized mean ΔF/F decreased with reduced imaging power, indicating diminished neuronal activity caused by crosstalk (Figure 2d). Specifically, the mean ΔF/F value at 15 mW excitation is more than double that at 5 mW. To further quantify neuronal activity, we applied the CNMF algorithm to deconvolve calcium traces and estimate “spike values”, which approximate spiking frequency^45^ (see Methods and Figure S5). Compared to 15 mW excitation, neuronal activity (spike values) was significantly reduced at 5 mW and 10 mW excitation (Figure 2g). On the contrary, in control experiments using ChR2-larvae (*Tg(KalTA4u508; UAS-GCaMP6s)*), neither mean ΔF/F nor spike values in KalTA4u508+ neurons varied across three excitation power conditions (Figure 2f, 2i). Notably, under 5 mW excitation, both metrics were statistically indistinguishable between ChR2+ and opsin-neurons (Figure S6), confirming the absence of detectable crosstalk under low excitation power. This finding is crucial for the effectiveness of the APPC method, as it selectively attenuates laser intensity in ChR2+ neurons to minimize crosstalk while maintaining the optogenetically stimulated calcium signal from ChR2+ neurons at an acceptable SNR.

**Figure 2.**
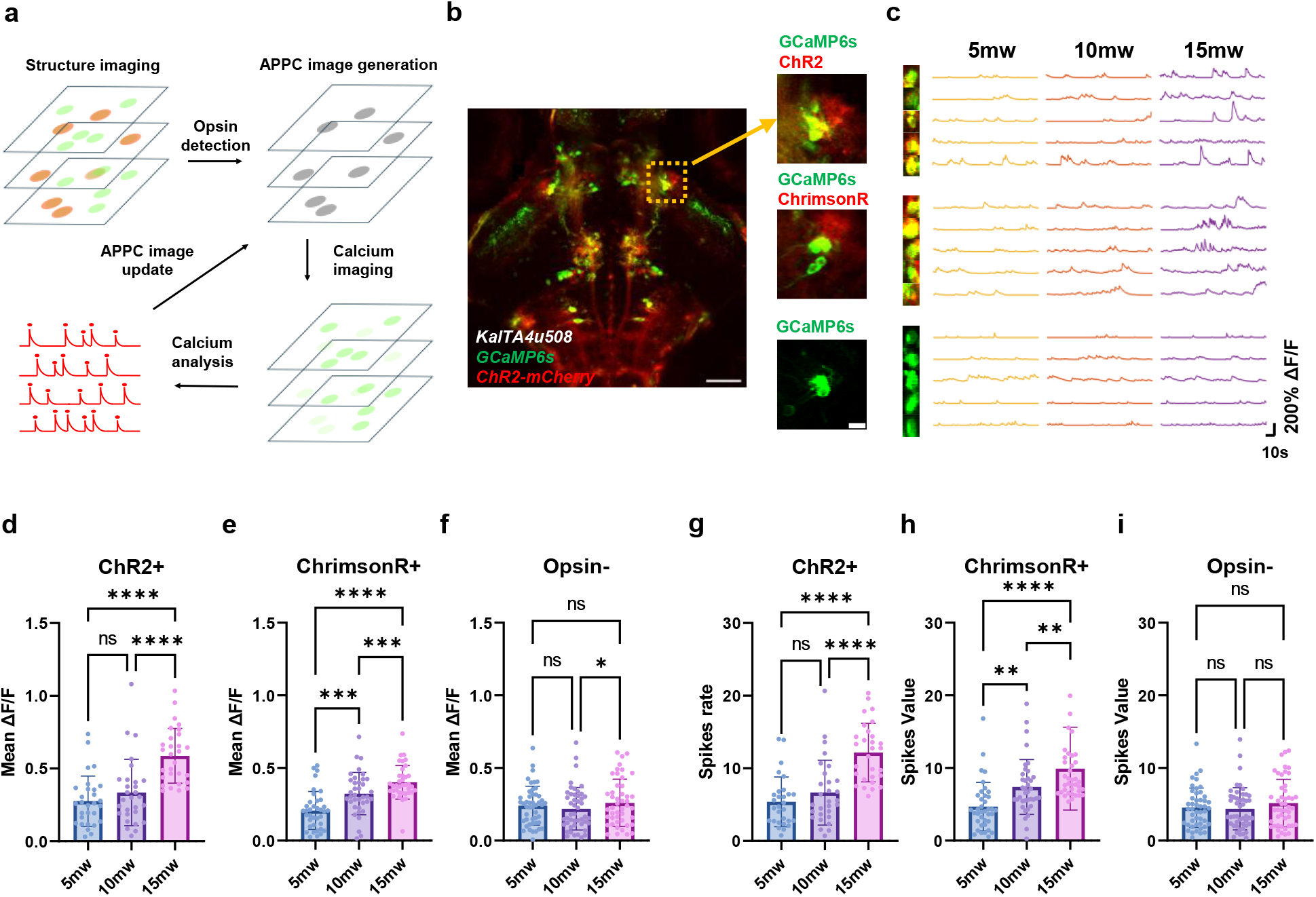
Active pixel power control minimizing crosstalk on ChR+ neurons. **a**, Experimental strategy for minimizing crosstalk of optical interrogation of ensembles. **b**, Left: Maximum intensity projection of a two-photon image stack encompassing the midbrain of a 7 dpf double transgenic *Tg(KalTA4u508;UAS:GCaMP6s; UAS:ChR2(H134R)-mCherry).* Right top: enlarged ROI for crosstalk evaluation. Right middle: enlarged ROI for crosstalk evaluation of 7 dpf double transgenic *Tg(KalTA4u508;UAS:GCaMP6s; UAS:ChrimsonR-mKate).* Right bottom: ROI of control group, *Tg(KalTA4u508;UAS:GCaMP6s).* **c**, Calcium signal recorded from GCaMP and with and without channelrhodopsin co-expressed neurons with different imaging power. Top group: ChR2+ neurons, middle group: ChrimsonR+ neurons, bottom group (control): ChR-neurons. **d**, Comparison of mean calcium fluctuation (mean±s.d.) of ChR2+ group under different image power (n=29 neurons, 7 fish, Friedman test with Dunn’s multiple comparisons test, 5 mW: 0.27±0.17, 10 mW: 0.33±0.23, 15 mW: 0.59±0.19, P=0.2635, ****P < 0.0001). e same as d but for ChrimsonR+ group (n=37 neurons, 8 fish, one-way analysis of variance (ANOVA) with Holm-Šídák test, 5 mW: 0.21±0.13, 10 mW: 0.32±0.15, 15 mW: 0.40±0.12, ***P = 0.0001 (5 mW vs 10 mW) and = 0.0009 (10 mW vs 15 mW), ****P < 0.0001) and, f same as d but for Opsin-group (n=47 neurons, 10 fish, Friedman test with Dunn’s multiple comparisons test, 5 mW: 0.24±0.13, 10 mW: 0.22±0.15, 15 mW: 0.26±0.16, P=0.2967 (5 mW vs 10 mW) and 0.9070 (5 mW vs 15 mW), *P = 0.0220). g, Spikes value (mean±s.d.) of ChR2+ group (n=29 neurons, 7 fish, Friedman test with Dunn’s multiple comparisons test, 5 mW: 5.38±3.42, 10 mW: 6.65±4.46, 15 mW: 12.15±4.03, P=0.4459, ****P < 0.0001) under different image power. h same as g but for ChrimsonR+ group (n=37 neurons, 8 fish, Friedman test with Dunn’s multiple comparisons test, 5 mW: 4.74±3.34, 10 mW: 7.08±3.30, 15 mW: 9.18±3.44, **P=0.0033 (10 mW vs 15 mW) and 0.0075 (10 mW vs 15 mW), ****P < 0.0001) and, **i** same as **g** but for Opsin-group (n=47 neurons, 10 fish, Friedman test with Dunn’s multiple comparisons, 5 mW: 4.54±2.71, 10 mW: 4.39±2.89, 15 mW: 5.16±3.28, P > 0.9999 (5 mW vs 10 mW), P=0.2386 (10 mW vs 15 mW) and 0.0698 (10 mW vs 15 mW)). Scale bar in **e** left is 50 μm and in **e** right is 10 μm.

ChrimsonR, a widely used red-shifted channelrhodopsin, is favored due to the separation of its single-photon excitation spectrum from that of GCaMP^46^. To test whether two-photon calcium imaging at 920 nm excitation would induce significant crosstalk in ChrimsonR-expressing neurons, we conducted a similar experiment in the *Tg(KalTA4u508; UAS-GCaMP6s; UAS-ChrimsonR-mKate2)* larvae^46^. The results showed that despite spectral separation of single-photon excitation, neurons expressing ChrimsonR exhibited similar behaviors of crosstalk as ChR2 for two-photon imaging: the reduced artefactual neuronal activity was observed under low-power excitation (Figure 2e). Spike values at 5 mW decreased significantly compared to 15 mW and 10 mW (Figure 2h), indicating the presence of residual crosstalk at high excitation power, which likely arises from ChrimsonR’s strong two-photon excitation at 920 nm (Figure S7). Again, both mean ΔF/F and spike values did not differ significantly between ChrimsonR+ and opsin-neurons under 5 mW illumination (Figure S6). Taken together, these results indicate that localized low-power imaging effectively mitigates crosstalk. It is important to note that ChrimsonR was fused with mKate2, which exhibits higher fluorescence intensity than the mCherry marker fused to ChR2^47^, thus complicating direct comparisons of channel expression levels based solely on fluorescent protein signals. A plausible explanation for the unexpectedly high crosstalk with ChrimsonR may be attributed to the elevated membrane expression of ChrimsonR, combined with sufficient two-photon excitation efficiency at 920 nm, collectively leading to the substantial crosstalk observed in our ChrimsonR-expressing neurons^6^.

In summary, we determined that 5 mW is a nearly crosstalk-free power for both ChR2 and ChrimsonR under our experimental conditions. While the calcium traces of ChR+ neurons could be reliably extracted due to their strong responses to optogenetic stimulation (Figure S4), downstream neurons typically exhibit significantly weaker or distinct responses compared to upstream neurons (Figure S8). Under low signal-to-noise ratio (SNR) conditions, downstream neurons with weak responses cannot be reliably identified or analyzed using the CNMF algorithm^45^(Figure S9). Therefore, it is essential to set the laser power in ChR-regions to 15 mW to ensure the quality of calcium signal data, which is crucial for accurately dissecting neural circuits. This demonstrates that APPC is critical for minimizing crosstalk while ensuring an acceptable SNR for calcium imaging of all neurons in the circuit.

### Interrogation of neural circuits with minimized crosstalk via active pixel power control

We first investigated how crosstalk at the ChR+ neurons affects their downstream neurons. To identify these downstream neurons, we applied two-photon holographic optogenetic stimulation targeted at pretectal u508+ neurons while simultaneously performing volumetric whole-brain imaging. Concurrently, we utilized APPC on the target u508+ neurons to control laser power for subsequent crosstalk evaluation (Figure 3a). In these experiments, we systematically reduced the imaging power at the ChR+ pixels from 15 mW to 10 mW, 5 mW, and “0 mW,” while maintaining 15 mW imaging power at other pixels to ensure high-quality calcium signals from downstream neurons. For this, we used larvae pan-neuronally expressing nucleus-localized GCaMP6s, while restricting ChR2 or ChrimsonR expression to the u508+ neurons Tg(*elavl3:Hsa.H2B-GCaMP6s*;*KalTA4u508; UAS:ChR2 (H134R)-mCherry*)^9,43,48^ or Tg(*elavl3:Hsa.H2B-GCaMP6s; KalTA4u508; UAS:ChrimsonR-mKate2*)^46^. We demonstrated that 920 nm optogenetic stimulation effectively activated neurons expressing both ChR2 and ChrimsonR (Figure 3b). However, we observed that the colocalization of GCaMP6s and ChRs was very low in both strains of zebrafish larvae. Consequently, our study focused on analyzing the calcium responses from downstream neurons.

**Figure 3.**
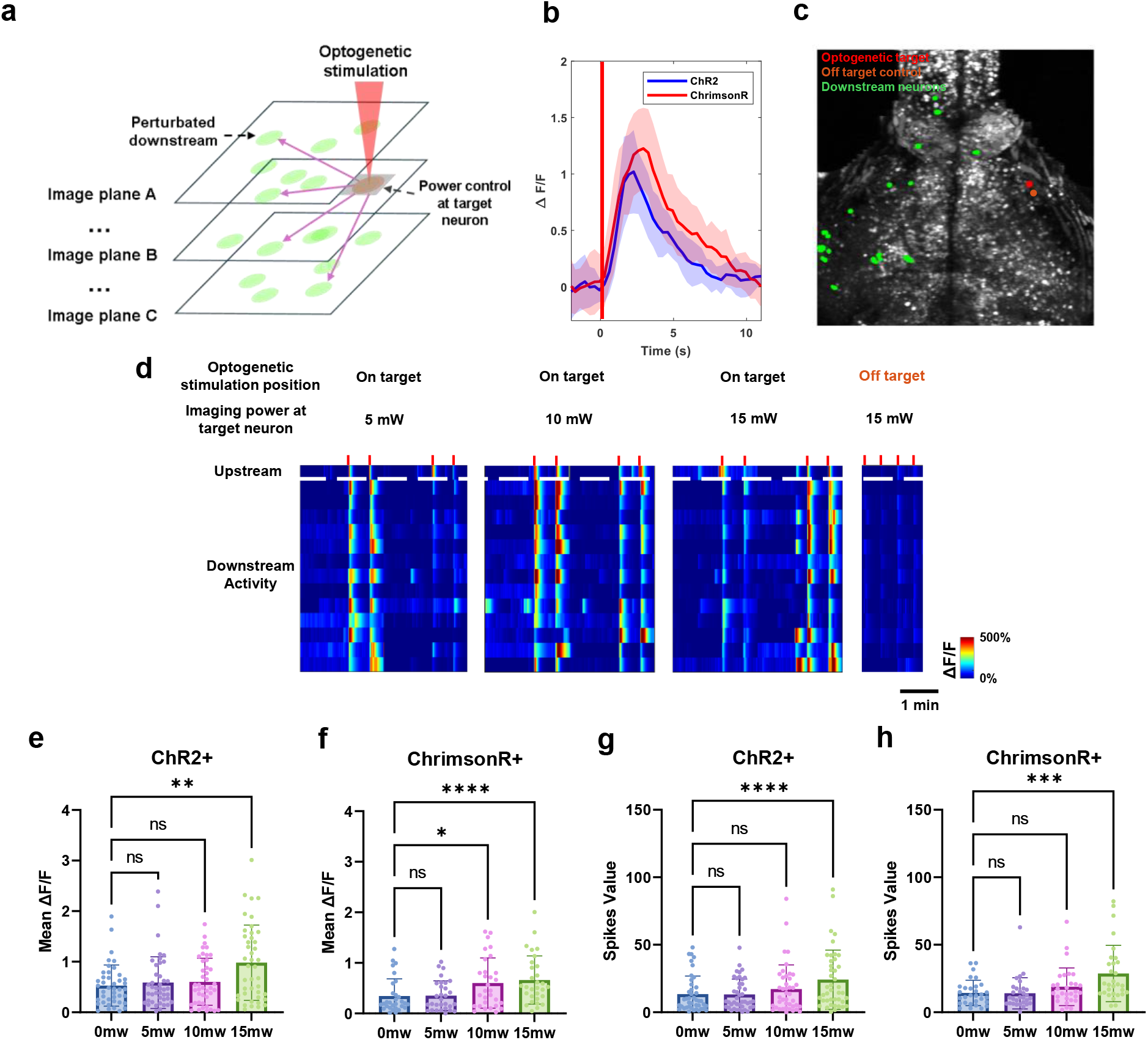
Active pixel power control mitigates crosstalk in neural circuits. **a**, Experimental strategy for evaluating crosstalk spreading in the downstream neurons. **b**, 920 nm two-photon optogenetics stimulation effectively activates both ChR2 and ChrimsonR. The somatic GCaMP signals (mean±s.d.) averaged across ten trails under the same optogenetic, with onset indicated by the red bar for 1 second. **c.** Field of view of GCaMP6s fluorescent signal of zebrafish brain depth projection across 80-180 µm. Red patch: optogenetic stimulation target. Orange patch: off-target control stimulation position. Green: putative downstream neurons. **d**, Heatmap displaying the activities of the optogenetic target neuron and 13 downstream neurons. The optogenetic target neuron was imaged under 5 mW, 10 mW and 15 mW, while the downstream neurons are imaged under 15 mW. The downstream neurons are determined based on the correlation between the regressor related to optogenetics stimulation (stimulation onset at red lines). **e,** Comparison of mean calcium fluctuation (mean±s.d.) of ChR2+ group downstream neurons under different image power on upstream neuron (n=42 neurons, 8 fish, Friedman test with Dunn’s multiple comparisons test, no optogenetic stimulation: 0 mW: 0.53±0.41, 5 mW: 0.59±0.51, 10 mW: 0.60±0.47, 15 mW: 0.98±0.74, P > 0.9999 (0 mW vs 10 mW), P = 0.8157 (0 mW vs 15 mW), **P=0.0022), **f,** same as **e** but for ChrimsonR+ group (n=30 neurons, 7 fish, Friedman test with Dunn’s multiple comparisons test, no optogenetic stimulation: 0 mW: 0.34±0.34, 5 mW: 0.35±0.29, 10 mW: 0.60±0.50, 15 mW: 0.66±0.48, P > 0.9999 (0 mW vs 5 mW) and *P=0.0153 (0 mW vs 10 mW), ****P <0.0001(0 mW vs 15 mW)). **g**, Spikes value (mean±s.d.) of ChR2+ group downstream neurons under different image power on upstream neuron (n=42 neurons, 8 fish, Friedman test with Dunn’s multiple comparisons test, no optogenetic stimulation: 0 mW: 13.40±13.47, 5 mW: 13.18±11.43, 10 mW: 17.21±17.86, 15 mW: 24.13±21.93, P > 0.9999 (0 mW vs 5 mW) and P=0.9315 (0 mW vs 10 mW), ****P < 0.0001(0 mW vs 15 mW)). **h**, same as **g** but for ChrimsonR+ group (n=30 neurons, 7 fish, Friedman test with Dunn’s multiple comparisons test, no optogenetic stimulation: 0 mW: 14.31±9.40, 5 mW: 14.00±11.59, 10 mW: 18.89±13.95, 15 mW: 28.67±20.86, P > 0.9999 (0 mW vs 5 mW) and P=0.8140(0 mW vs 10 mW), ***P = 0.0010(0 mW vs 15 mW)).

The putative downstream neurons were identified by their highly correlated activity patterns with the stimulated u508+ neurons^49^. Since the stimulation light can scatter onto the adjacent retina, potentially causing visual input artifacts, we conducted control experiments to exclude sensory neurons responding to this scattered light. Specifically, we positioned the optogenetic stimulation light in a neighboring region of the ChR+ neurons (Figure 3c). Only neurons that responded specifically to pretectal ChR activation, and not generally to light (Figure S10), were selected as downstream neurons (Figure 3d). These neurons exhibited strong correlations with the optogenetic stimulation applied to the targeted ChR+ neuron across various imaging conditions (Figure 3e). In zebrafish expressing either ChR2 or ChrimsonR in the u508+ neurons, analysis of the downstream neurons revealed that their artefactual activity diminished as the imaging power applied to the upstream neurons was reduced (Figure 3e-h). Importantly, these results demonstrate that applying 15 mW imaging power to ChR+ regions caused strong crosstalk, which affected downstream neurons and perturbed the neuronal circuit. Furthermore, neuronal activity levels were statistically indistinguishable when the upstream neurons were scanned at 5 mW versus “0 mW” (Figure 3e-h). This led to the conclusion that 5 mW is the “crosstalk-free” power for upstream neurons, effectively minimizing crosstalk and providing a solution for negligible interference within the neuronal circuit. This means that in a strain of zebrafish expressing calcium indicators and ChRs in upstream neurons, alongside calcium indicators in downstream neurons, crosstalk-free optical interrogation can be achieved using the APPC approach.

To extend our method to different imaging scenarios, we investigated how to optimize imaging conditions to minimize crosstalk. A particularly important consideration is the extent to which ChRs can be activated by out-of-focus scanning light in 3D volumetric imaging. First, we modeled the activation of ChR2 molecules during imaging (see Methods) and assessed the extent of ChR2 molecule activation under various scanning conditions. Our simulation indicates that the ChR2 molecule can be effectively activated at a typical low power density of ∼0.2 mW/µm^2^ with a stimulation duration of 1 ms, consistent with numerous *in vivo* studies^5,6,35,50^ (Figure 4a and Figure S11). The excitation probability is proportional to laser power density and stimulation time. However, during calcium imaging, the power density at the focal point can be 50 times higher than the typical activation power density, significantly reducing the stimulation time required for ChR activation^1,9,51^. Under 15 mW excitation, a ChR2 molecule has an activation probability of ∼8.1%, which drops to ∼0.9% at 5 mW excitation, under a pixel dwelling time of 68 ns corresponding to our imaging conditions (Figure 4a). The trend is consistent with our previous *in vivo* experiment results. In addition, a recent study also demonstrated that rapid raster scanning can lead to power-dependent crosstalk^52^. Therefore, by controlling imaging power at the pixel level (e.g., 5 mW), we can effectively mitigate crosstalk. Suitable imaging power can be further adjusted according to the change of FOV. A previous study has shown that setting stationary out-of-focus light with a high numerical aperture (NA) can activate more ChRs across the entire membrane compared to focusing on the soma equator^34^. To simulate the scanning scenario, we set the neuronal soma diameter to 10 µm and assumed that ChR2 expression is confined to the soma membrane. The imaging light was scanned above or below the soma equator at varying distances and power levels (Figure 4b). Our simulation indicated that the total photocurrent decreases as the distance between the focal plane and the equator increases (Figure 4c and Figure S12). Given the symmetry of the photocurrent generated by the out-of-focus plane, we focused only on the scenarios where the out-of-focus plane was positioned above the equator. To validate our simulation results *in vivo*, we imaged on ChR2+ neurons in larva zebrafish *(Tg(KalTAu508; UAS-ChR2(H134R)-mCherry; UAS-GCaMP6s)* ^48^ by positioning one imaging plane at the soma equator with a power of 5 mW for calcium recording, while moving the position of the second imaging plane (under 15 mW excitation) from 0 µm to 14 µm, exceeding the typical sampling gap^9,42^. Consistent with our simulation results, out-of-focus light at 10 µm or farther away from the equator leads to a negligible crosstalk (Figure 4d). These results provide critical guidance for determining optimal depth intervals in 3D volumetric imaging protocols.

**Figure 4.**
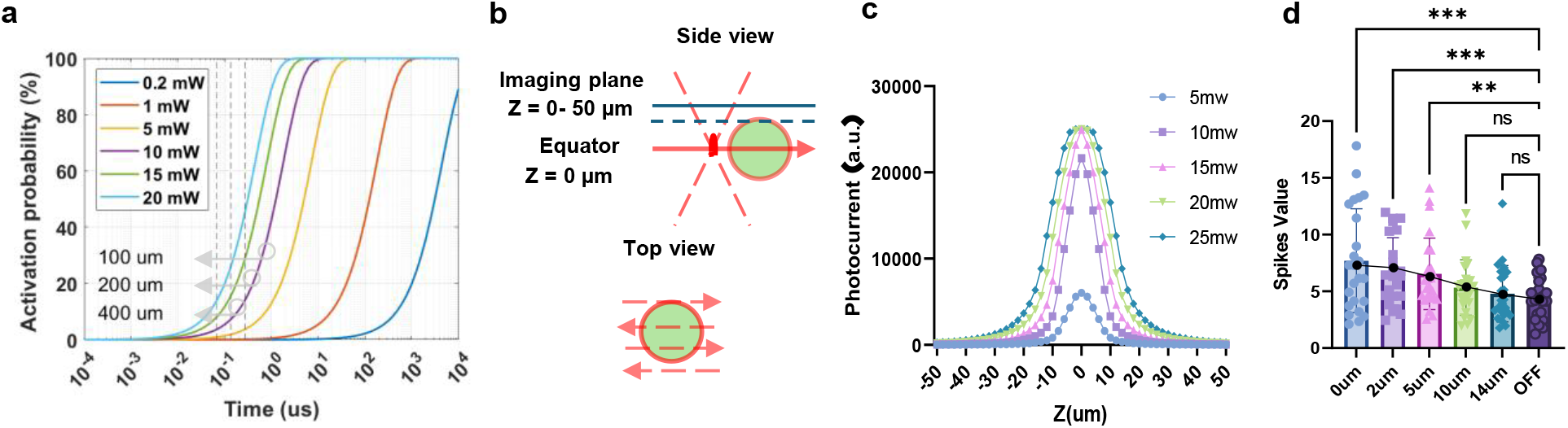
Activation of ChR2 under different stimulation conditions. **a,** The activation probability of a ChR2 molecule under stimulation by two-photon raster scanning at different powers of a defocused laser (Simulated N.A. = 0.9). The dashed lines correspond to the stimulation times for 100 μm, 200 μm, and 400 μm FOVs using a resonant scanner with an 8 kHz resonance frequency. The results account for the attenuation caused by the sample tissue (see Methods). For simulation results that do not consider attenuation for *in vitro* experiment as reference, see Figure S11. **b**, Illustration of raster scanning through the entire focal plane at a distance z from the cell equator. Green circle: soma. Red line: laser scanning path. **c**, Simulated photocurrents stimulated by raster scanning at various distances Z related to the equator. **d**, The spikes value (mean±s.d.) of target neuron with an extra scanning plane outside of cell equator (n=24 neurons, 15 fish, Friedman test with Dunn’s multiple comparisons test, 0 um: 7.68±4.57, 2 um: 6.75±2.97, 5 um: 6.54±3.16, 10 um: 5.33±2.44, 14 um: 4.79±2.48, 10 um: 4.35±1.85, ***P =0.0002 (0 um vs OFF), 0.0004(2 um vs OFF), **P=0.0019, P=0.4482 (10 um vs OFF) and P>0.9999 (14 um vs OFF)). Black dot and black line: simulated results in **c** and fitted to spike value.

## Discussion

In summary, we have developed the APPC approach, a sophisticated all-optical methodology, for simultaneous neuronal activity monitoring and manipulation while substantially reducing crosstalk interference. By exploiting the rapid tuning capabilities of AOM, we achieved continuous modulation of scanning light intensity with single-pixel resolution. Based on ChR expression patterns, we implemented dynamic control of imaging power within precisely defined regions of interest to prevent inadvertent ChR activation by imaging light. In larval zebrafish experiments, our methodology facilitated comprehensive whole-brain functional imaging while mitigating laser-induced behavioral artifacts through selective inhibition of scanning in the ocular region. This approach demonstrates significant potential for preserving the functional integrity of multiple light-sensitive domains within the imaging volume. Furthermore, we established that a single femtosecond laser can effectively perform concurrent volumetric calcium imaging and two-photon holographic optogenetic stimulation, even when utilizing red-shifted ChrimsonR.

Our investigation initially demonstrated that both ChR2 and ChrimsonR exhibit significant crosstalk when exposed to conventional excitation laser power, potentially modulating downstream neuronal dynamics. This finding indicates that laser scanning not only directly affects ChR+ neurons but also disrupts broader neuronal circuit functionality. The magnitude of crosstalk varies depending on ChR expression levels and intrinsic properties (such as excitation spectrum, kinetics, and photocurrent)^1,14,27^, which were analyzed through rigorous electrophysiological methods^1,4,6,26,51,52^. In this study, we present a comprehensive framework for the optical estimation of crosstalk and the adaptive optimization of imaging conditions. Specifically, our approach dynamically adjusts the optimal imaging power for each target neuron based on factors such as location (imaging depth), ChR expression levels, and the intrinsic properties of neurons. This enables the minimization of crosstalk while maintaining high-fidelity calcium signal readout. Additionally, we developed a computational simulation model to predict viable imaging parameters. By integrating optical neuronal activity assessments with dynamic excitation power control, we identified “crosstalk-free” imaging conditions that minimize interference for both upstream and downstream neurons. These conditions were validated through computational simulations and *in vivo* volumetric imaging. Furthermore, our methodology allows for the selective amplification of laser power in specific regions to enhance the signal-to-noise ratio while reducing thermal effects on biological specimens.

## Methods

### Animals

All procedures conformed to the guidelines by the animal ethics committee of HKUST. Animals were kept under a standard 14:10 light cycle at 28°C. Zebrafish larvae carrying mutations in the *mitfa* allele (nacre) were used for all experiments. Larvae were kept in Danieau’s solution, and fed with paramecia (*Paramecium multimicronucleatum*, Carolina Biological Supply Company) on 5 to 7 days post fertilization (dpf).

The following transgenic lines were used for this study: *Tg(KalTA4u508;UAS:GCaMP6s; UAS:ChR2(H134R)-mCherry)^44,50^, Tg(KalTA4u508;UAS:GCaMP6s) ^43^, Tg(KalTA4u508; UAS:GCaMP6s; UAS:ChrimsonR-mKate2)^46^, Tg(elavl3:Hsa.H2B-GCaMP6s; KalTA4u508; UAS:ChR2(H134R)-mCherry), and Tg(elavl3:Hsa.H2B-GCaMP6s;UAS:ChrimsonR-mKate2)^48^*.

### Zebrafish preparation

6-8 dpf larvae were embedded in 2% low melt agarose (Invitrogen) on 90 mm round (Thermo Scientific (cat#101VR20)) plastic petri dish. The agarose surrounding the tail and the eyes was removed by surgical blade (Paragon p303) to allow for tail and eye movements. Fish were allowed to recover overnight prior to the experiment.

### Two-photon imaging microscope

A home-built two-photon light scanning microscope controlled by a C# based custom software is modified from previous setup. The laser source is a Ti:Sapphire femtosecond laser (Chameleon Ultra II, Coherent), repetition rate: 80 Mhz, and can be tuned between 680 nm and 1080 nm. The laser beam first passes through the half-wave plate to adjust polarization direction. Then, a polarized beam-splitter separates the laser beam into two paths, one for imaging and the other for optogenetic stimulation.

At imaging path, the laser beam first pass through a 4f system of scale factor x0.467 (AC254-075-AB-ML F = 75 mm, Thorlabs and AC254-035-AB-ML F = 35 mm, Thorlabs) in order to not overfill the AOM (CAOM-080-010-TEC-740-1020-AF-A17, Castech) aperture (1.0mm). After passing through the AOM, the laser beam is expanded by a factor of ×3 through a 4f system (AC254-050-AB-ML F = 50 mm, Thorlabs and AC254-150-AB-ML F = 150 mm, Thorlabs). Two dispersion compensating prism (SF10 25.4 x 25.4mm Ultrafast Prism, Edmund optics) is set and optimized the dispersion compensation at 920nm. The first prism is fixed, and the second prism is located at a linear translation stage with actuator (CONEX-LTA-HL, Newport). The working direction of the linear translation stage is perpendicular to the prism base plane. For each wavelength used, the position of the prism is adjusted by maximizing the fluorescent intensity from the fluorescent plate. The total system group dispersion was estimated from the optical components used to calculate the distance between two prisms when the laser worked at 920 nm. The pulse duration after the objective is 118 fs, measured by pulseCheck autocorrelators (NX 150, A.P.E.) to verify the dispersion compensation.

The laser beam with 4 mm in diameter (propagate a large distance and thus expanded) is then expanded by a factor of ×4 through a 4f system (AC254-050-AB-ML F = 50 mm, Thorlabs and AC254-200-AB-ML F = 200 mm, Thorlabs) to full fill the ETL (EL-16-40-TC-NIR, Optotune AG), which is used for multi-plane functional imaging. Then a 4f system (AC254-200-AB-ML F = 200 mm, Thorlabs, and AC254-050-AB-ML F = 50 mm, Thorlabs) reduces the beam diameter to 4 mm to avoid overfilling the scanner mirror. A resonant scanner (CRS 8 kHz, Cambridge technology) for high-speed scanning is conjugated with a set of XY galvo scanners (6215H, Cambridge technology) by a 1:1 relay (AC254-100-AB-ML F = 100 mm, Thorlabs). Finally, two lenses (Scan lens SL50-CLS2 F = 50 mm, Thorlabs, and Tube lens TTL200MP F = 200 mm, Thorlabs) are used to correct the aberration and expand the laser beam to fulfill the objective back pupil (objective lens (20x, N.A. 1.0, XLUMPLFLN20XW, Olympus Corporation). The ETL, resonant scanner, XY galvo scanners, and objective pupil plane are all conjugated. In the fluorescence collection path, a dichroic mirror (FF705-Di01-25*36, Semrock) is set between the tube lens and objective to reflect the fluorescence light, which then passes through two relay lenses (AC254-150-AB-ML F = 150 mm, Thorlabs and AC254-040-AB-ML F = 40 mm, Thorlabs) into the PMT (H11461-03, Hamamatsu). Two filters (FF03-525/50-25, Semrock, and FF01-600/52-25, Semrock) are installed on a flipper which can switch the filter to collect different fluorescence signals. Voltage signals are converted from PMT output current using a high-speed current amplifier (DHPCA-100, Femto) and digitized using an oscilloscope device (PCIe-8512H, ART technology).

At stimulation path, the laser beam passes through a set of galvo scanners (6215H, Cambridge Technology). Then the light beam is expended by two lenes (AC254-50-AB-ML F = 50 mm, Thorlabs, and AC254-250-AB-ML F = 250 mm, Thorlabs) to fill the spatial light modulator (HSP1K-500-1200-PC8, Meadowlark Optics). Then the SLM is conjugated to the back focal plane of objective by a demagnify relay (AC508-250-AB-ML F = 250 mm, Thorlabs, and AC254-150-AB-ML F =150 mm, Thorlabs). The photon stimulation light combines with imaging light by a polarized beam splitter (CCM1-PBS252/M; Thorlabs) between tube lens and dichroic mirror.

The LCR projector and camera were employed to provide visual stimuli and record animal behaviour, as detailed described in our previous work^49^.

### Pixelwise adjustment of laser intensity

The orientation of the AOM is initially adjusted to achieve maximum diffraction efficiency. Here we adopt the 0^th^ order light and measured the intensity modulation dynamic range, which spans from 5% to 100%.

To enable dynamic adjustment of laser intensity on a pixel-by-pixel basis, we developed a GUI that allows for the custom generation of figures using fluorescence images of ChR+ neurons as the pattern for APPC. These figures are encoded into voltage signals and transmitted to the AOM driver (CARD-FE2-080-24D04C-1DA5H-AF, Castech). To synchronize the AOM with the imaging scanning beam, the clock signal generated by the resonant scanner serves as the reference clock signal for the DAQ card (PCIe9105_H, ART technology). The AO port is configured to generate a series of output signals upon receiving a trigger from the resonant scanner clock, thereby eliminating time delay accumulation (Figure S3). Finally, we captured the fluorescence image from a fluorescence plate with the loaded figure to finely tune the delay by aligning the fluorescence pattern between two adjacent lines.

### *In vivo* calcium imaging and holographic optogenetics stimulation

Two-photon multiple-plane imaging was performed using resonant-galvanometer raster scanning (30 Hz frame rate, 512x512 pixel per frame, 400 µm field-of-view, 12-14 µm between imaging plane, 10 planes, 3 Hz volume rate) typical for whole brain imaging in zebrafish^49^. An objective lens (20x, N.A. 1.0, XLUMPLFLN20XW, Olympus Corporation) was used for all experiments. The femtosecond laser (Chameleon Ultra II, Coherent) was operated at 920 nm wavelength for calcium imaging and 1020 nm for mCherry and mKate imaging. In all the *in vivo* experiments, the imaging power applied to the sample ranged from 0.5 to 15 mW. At the beginning of the imaging, we first imaging at the very low power, 0.5 mW, to outline the eye contours from the fluorescence signal of the pigments on retina. Then we manually selected the eye region and included it in the APPC pattern, where the laser power was set to be the minimum. For pixel level power control of ChR expression region, the APPC pattern was generated based on the ChR expression pattern verified by fused fluorescent protein. We set the pixel intensity threshold to adjust the pattern region flexibility. We then manually refined the pattern for each plane. The imaging power at these regions were determined based on the experiment requirement.

For optogenetics stimulation, the holograms were calculated by compressive sensing weighted Gerchberg–Saxton algorithm^53^ and loaded on SLM which controlled by the custom software in MATLAB. A large beam diameter is applied to achieve an effective excitation NA (∼0.5) for axial profile compression. A spot array is then created on each target neuron, consisting of spots that are 10-12 µm in diameter with a 1 µm gap between them, and an average power density of 0.15-0.3 mW/µm^2^. The spot array is scanned over a range of 2 µm to ensure that the entire soma area is stimulated. To mimic natural neuronal response, we first provided visual stimuli and extract the calcium signal from the anterior pretectal u508 neurons. Then we adjust the optogenetic stimulation laser power to trigger similar calcium response. Each optogenetic stimulation lasted for 1 second, with a total of 5 stimulations per session, separated by 30-second intervals. In the experiments that aimed at identifying the downstream neurons, we selected approximately 5-10 anterior pretectal u508 neurons on either the left or right side.

Before each optogenetic experiment, we calibrated the alignment between the imaging light and the photo-stimulation light. We bleach two dark spots on a fluorescent microscope slide (FSK3, Thorlabs) using both imaging light and photo-stimulation light. After measuring the distance between the spots, we included a coordination shift in the hologram calculation.

### Calcium imaging data processing

Image analysis was performed using custom-written scripts in Matlab (Mathworks). Non-rigid motion correction for each imaging plane was performed using the NoRMCorre algorithm^54^. For neuron-based analyses, the locations of regions of interest (ROIs) and their fluorescent signal trace were extracted using the CNMF algorithm from the CaImAn package rate^45^. Image sections with axial shift > 5 µm were discarded because they were not able to keep precise information. ROIs with the SNR, as computed by CaImAn, below 1.5 were excluded from further analysis. Specifically, the deconvolveCa function returned the baseline, denoised calcium trace, noise level estimated from the spectral density and deconvolved signal (spikes train). We calculated ΔF/F based on the inferred baseline and calcium trace. The mean ΔF/F is calculated as the average ΔF/F value across all frames during the entire video recording. Then we computed the inferred spike value as the summation of the spike value normalized by the baseline. Spike values below one standard deviation of the noise level were discarded.

### Data presentation and statistical analyses

All values are presented as mean ± standard deviation (s.d.), unless otherwise specified. Statistical analyses were conducted using appropriate tests, as indicated in the figure legends. These statistical tests were performed using Graph Pad Prism, version 9.5.1, with a p-value of <.05 considered statistically significant. Data normality was first checked using the Shapiro–Wilk normality test. Normally distributed data were analyzed using paired, unpaired one-way ANOVA test. Non-normally distributed data was analyzed using paired, unpaired non-parametric test (Friedman test with Dunn’s multiple comparison test). In the statistical results presented in Figures 2d-i and 3e-h the neurons depicted were selected based on their distribution, as they exhibited strong crosstalk. Specifically, there was no statistically significant difference in spike values below laser power of 5 mW between the ChR+ and ChR-groups, leading us to consider crosstalk negligible at this power level. Neurons with mean ΔF/F or spike values under 15 mW imaging power exceeding the mean plus one standard deviation under 5 mW were classified as exhibiting strong crosstalk and included in Figures 2d-i and 3e-h.

A regressor for single optogenetic stimulation was generated by averaging the calcium trace corresponding to optogenetic stimulation. The downstream neurons depicted in Figures 3c-h were selected based on the criterion that the correlation between the calcium trace and the target optogenetic regressor exceeded 0.3^9^, while the correlation with the control optogenetic regressor remained below 0.05.

### ChRs modelling

A ChR2 molecule has an absorption rate of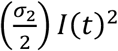, and the ground state lifetime is given by 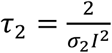, where *σ_2_* is the two-photon absorption cross-section (260GM)^34^. The laser intensity *I* which can be expressed by^55^:

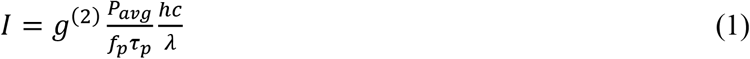

where *g*^(*2*)^ is the factor of 0.59 for a hyperbolic-secant-squared pulse shape^56^, *P_avg_* is the total time-average laser power, *f_p_* is the pulse train repetition rate, τ*_p_* is the pulse duration. In our calculations, we used *f_p_* = 80 *MHz* and τ*_p_* = 120 *fs. h* is the Planck’s constant, *c* is the light speed, *λ* is the wavelength.

The laser is attenuated in the sample due to scattering and absorption. We determined the attenuation length (AL) from the brain z-stack images, where the average fluorescence signal of the brightest 0.1% of pixels attenuates by 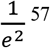 (Figure S13a). The calculated AL for 7-8 days post-fertilization (dpf) zebrafish larvae is approximately 201 µm (Figure S13b). At a depth of 140 µm, the laser power reaching the pretectal u508 neurons was estimated to be 50% of the initial power, based on the calculated AL, and this value was used for subsequent photocurrent calculations.

The probability of activating a ChR2 molecule under stationary illumination of time Δ*t* is:

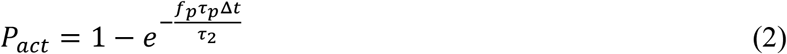

Next, we consider the activation of a ChR2 molecule by a moving focused beam, which has a Gaussian distribution in the lateral direction. Thus, the laser intensity can be expressed as^34^:

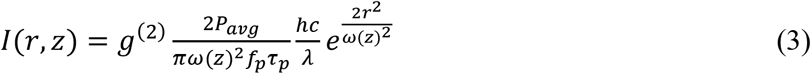

where *ω* is the beam waist. At the focal plane, *ω* = *ω*_0_ and *ω*_0_ can be determined by measuring the two-photon excitation point-spread function from image 0.2 um microspheres. The two-photon excitation waist *ω_TPE_* was obtained by fitting the two-photon excitation point-spread function by the Gaussian distribution. The *ω* can then be calculated from 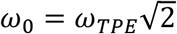. Or *ω*_0_ can be estimated by 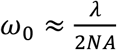. In the perifocal region, the *ω* can be expressed by^35^:

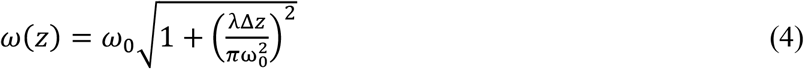

To simulate the activation of ChR2 molecules on the neuron, we modeled the neuronal soma as a sphere with a diameter of 10 µm, generating 25,000 points representing ChR2 molecules randomly distributed on the surface, corresponding to approximately 200 proteins proteins/µm^2^. We simulated the focused beam scanning and calculated the photocurrent of each ChR2 molecules by combining equ. (1)-(4) and integrating over time. The kinetic information can refer to Rickgauer’s work^34^. For ChR2, the photocycle can be completed before the next frame even for a 30Hz frame rate. Thus, we only considered the ChR2 activation during single frame scanning. The maximum photocurrent for each ChR2 molecule was normalized to 1. The total photocurrent for the entire neuron was then obtained by summing the photocurrent contributions from each individual ChR2 molecule.

## Acknowledgments

This work is supported by Hong Kong Research Grant Council to J.Y.Q. (16102122, 16102123, 16102421, 16102518, 16102920, T13-607/12R, T13-605/18W, C600217GF, C6001-19E, C6034-21G, T13-602/21N), the Innovation and Technology Commission (ITCPD/17-9), the Area of Excellence Scheme of the University Grants Committee (AoE/M-604/16, AOE/M-09/12) and the Hong Kong University of Science & Technology (HKUST) through grant 30 for 30 Research Initiative Scheme, and Hong Kong Research Grant Council to J.L.S. (16103522, 16101221, 16103118, and 26100617), State Key Laboratory of Molecular Neuroscience, The Hong Kong University of Science and Technology, Hong Kong, PR China, Innovation and Technology Commission (ITCPD/17-9).

## Author contributions

G.Y., G.T., and J.Y.Q. conceived the research idea and designed the experiments. G.Y. built the imaging systems. G.Y.; G.T. and K.Y.C performed animal surgery and imaging experiments under the supervision of J.Y.Q. and J.L.S.; Y.F. and Z.S. provided technical support; G.Y. and G.T. analyzed the data with assistance from H.Y.; G.Y., G.T., Y.H., J.L.S., and J.Y.Q. wrote the paper with input from all other authors.

